## Supplement Material for "*Akkermansia muciniphila* identified as key strain to alleviate gut barrier injury through Wnt signaling pathway"

Xinyan Han

College of Animal Sciences, Zhejiang University, 866 Yuhangtang Road, Hangzhou 310058, China

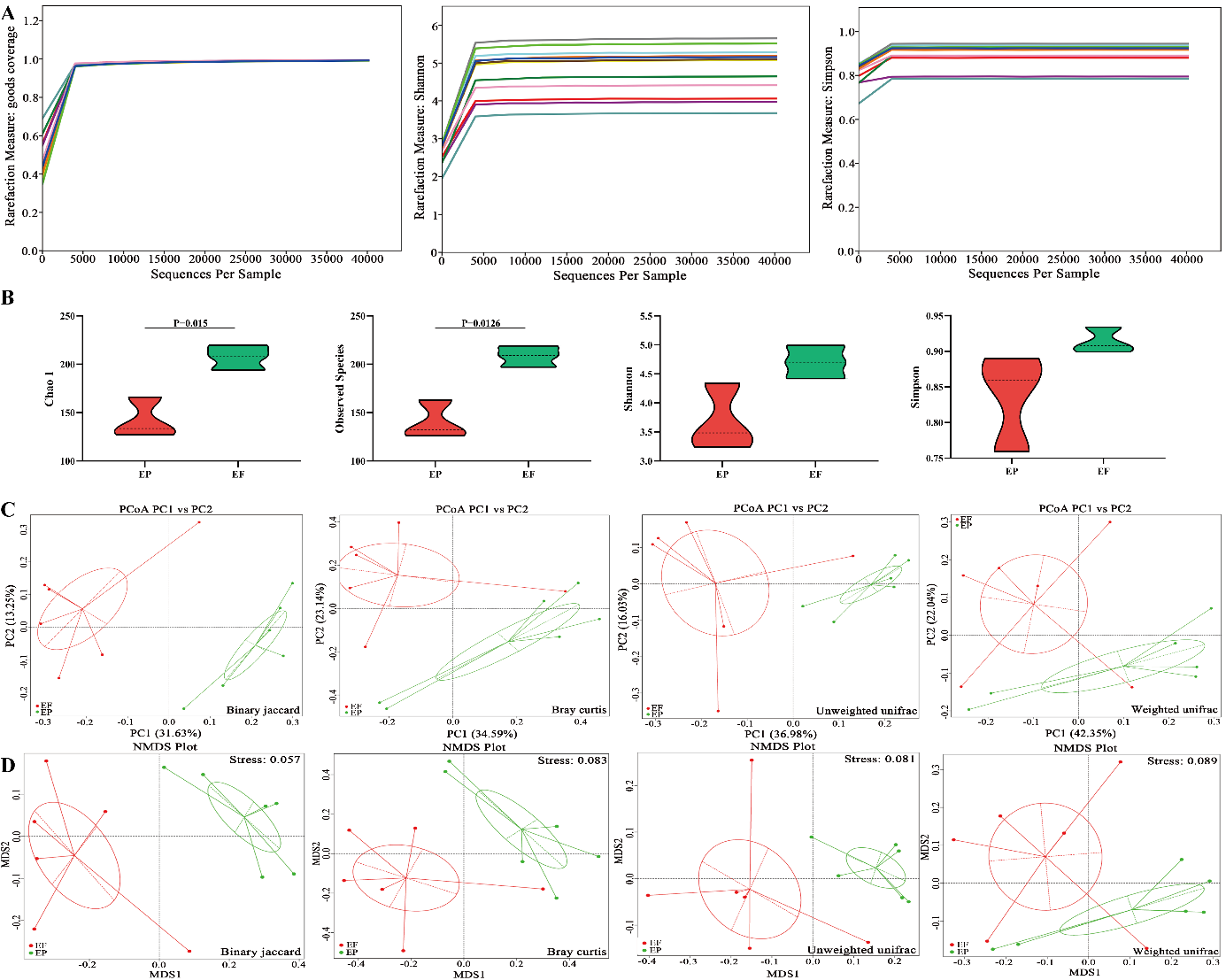

**Figure S1.** FMT altered the structure of gut microbiota in AIMD piglets infected with ETEC K88. (A) Rarefaction curves based on goods coverage, Shannon and Simpson index. (B) Alpha diversity indexes (Chao 1, Observed spices, Shannon and Simpson). (C) Principal coordinate analysis (PCoA) based on Binary jaccard, Bray curtis, Unweighted unifrac and Weighted unifrac. (D) Non-metric multidimensional scaling (NMDS) analysis based on Binary jaccard, Bray curtis, Unweighted unifrac and Weighted unifrac. EP: ETEC K88 + PBS group, EF: ETEC K88 + FMT group.

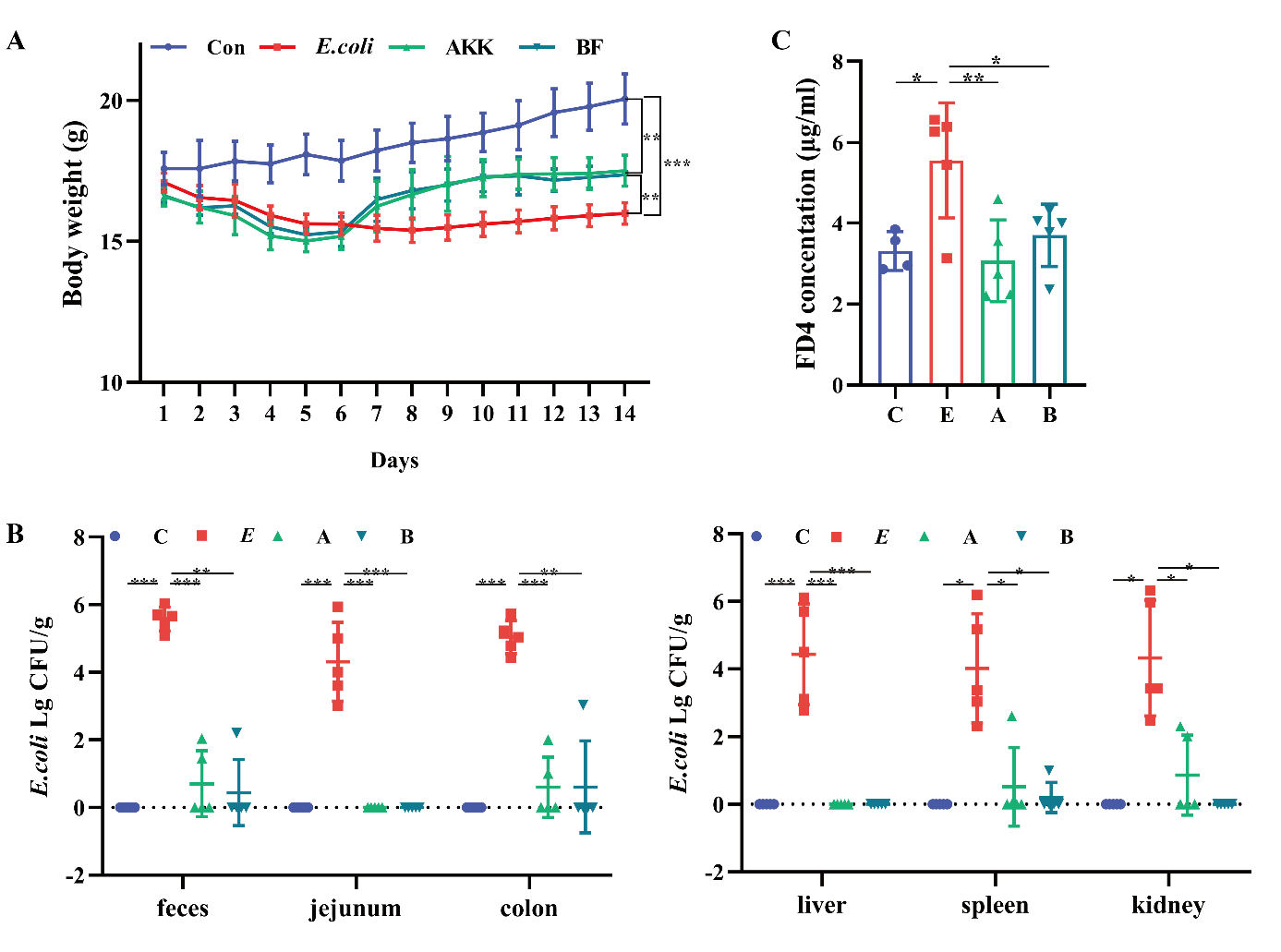

**Figure S2.** (A) The body weight of mice. n=6. (B) The *Escherichia coli* (*E. coli*) translocation in tissues and organs of mice. (C) The concentrations of serum 4 kDa fluorescein isothiocyanate-dextran (FD4) in C, E, A and B groups. C: control group, E: ETEC K88 + PBS group, A: ETEC K88 + *A. muciniphila* group, B: ETEC K88 + *B. fragilis* group. Data are expressed as the mean ± SD. *, *P* < 0.05, **, *P* < 0.01***, *P* < 0.001.

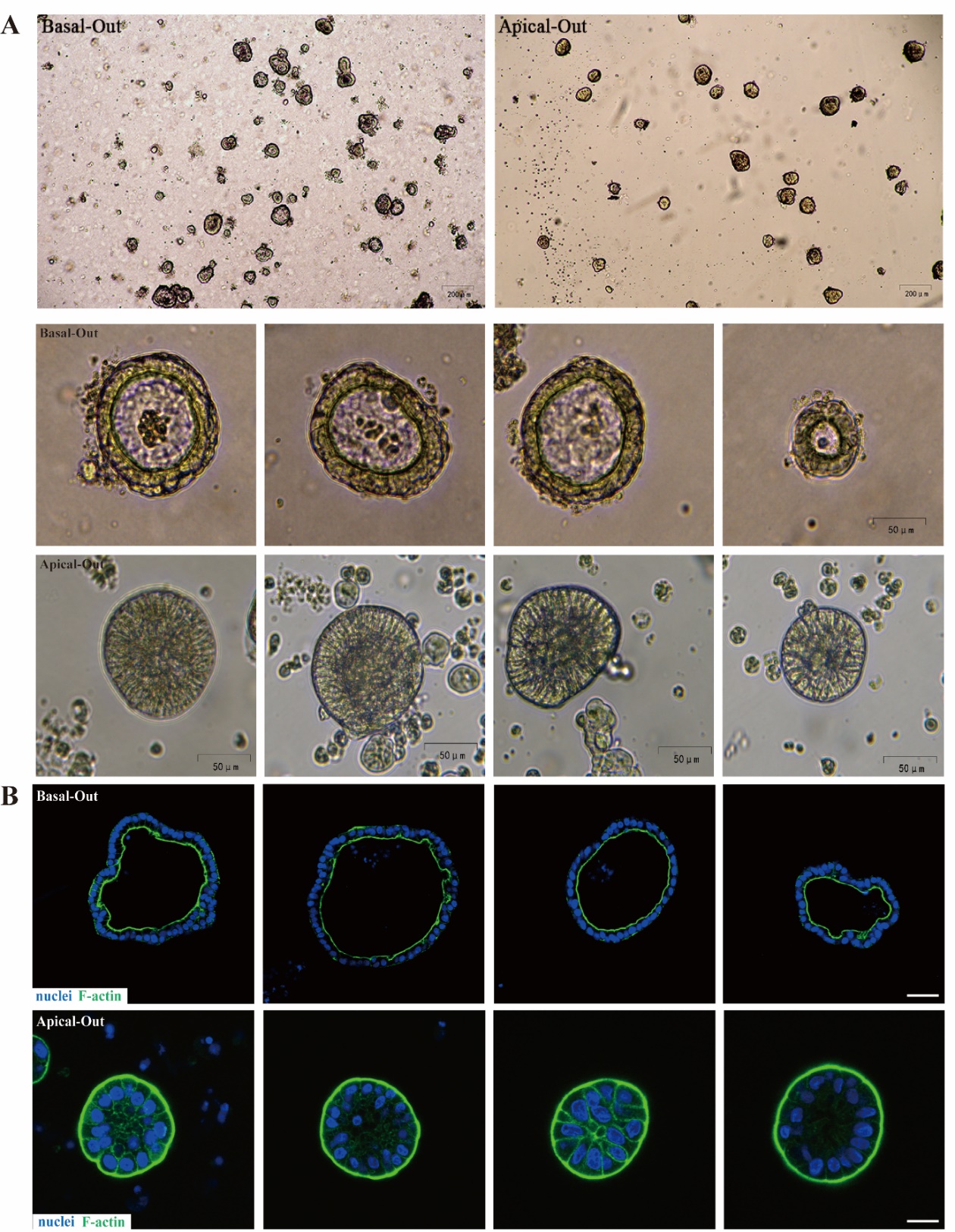

**Figure S3.** (A) Light microscope visualization of basal-out and apical-out organoids (4×, 20×). (B) Immunofluorescence images of F-actin in basal-out and apical-out intestinal organoids (scale bars = 20 μm).

**Table S1.** Real-time quantitative PCR primers and conditions.

| Gene | Primer sequences (5' to 3') | Genbank accession / Reference |
| --- | --- | --- |
| ETEC K88 | F: CGCCAGTAACTGGTGGTGTA | M25302.1 |
|  | R: CCATCAGGGTTTCTGAGTACTAC |  |
| Pig-  *IL-10* | F: GACCAGATGGGCGACTTGTTG | NM_214041.1 |
|  | R: GGGAGTTCACGTGCTCCTTGAT |  |
| Pig-  *TGF-β1* | F: GAAGCGCATCGAGGCCATTC | NM_214015 |
|  | R: GGCTCCGGTTCGACACTTTC |  |
| Pig-  *TNF-α* | F: CGCTCTTCTGCCTACTGCACTT | NM_214022.1 |
|  | R: CGGCTTTGACATTGGCTACAA |  |
| Pig-  *IL-1β* | F: GCCAGTCTACATTGCTCATGTTTCT | NM_001005149.1 |
|  | R: GTTGTCACCATTGTTAGCCATCAC |  |
| Pig-  *IL-6* | F: GCCTTCAGTCCAGTCGCCTTCT | NM_214399.1 |
|  | R: GTGGCATCACCTTTGGCATCTTC |  |
| Pig-  *IFN-γ* | F: CAGAGCCAAATTGTCTCCTTCTAC | NM_213948.1 |
|  | R: GTCATTCAGTTTCCCAGAGCTACCA |  |
| Pig-  *ZO-1* | F: GAGGCTCAGCCCTATCCATCTG | XM_021098856.1 |
|  | R: CGGGACCTGCTCATAACTTCGT |  |
| Pig-  *Occludin* | F: CGGCCATATCCAGAGTCTTCGT | NM_001163647.2 |
|  | R: CGTTTTGAAGACGCCTCCAAGT |  |
| Pig-  *Claudin1* | F: CCTACGCTGGTGACAACATTG | NM_001244539.1 |
|  | R: GTGGTGTTCAGATTCAGCAAGGA |  |
| Pig-  *E-Cadherin* | F: CCCCAACACTTCTCCCTTCACT | EU805482.1 |
|  | R: CTCGAGGGTTTTCTTTGGCTTC |  |
| Pig-  *β-catenin* | F: GCTGCTGTTTTGTTCCGAATGTC | NM_214367.1 |
|  | R: CCTGGGCACCAATATCAAGTCC |  |
| Pig-  *Lgr5* | F: CCTTGGCCCTGAACAAAATA | Gonzalez et al., 2013 |
|  | R: ATTTCTTTCCCAGGGAGTGG |  |
| Pig-  *Ki67* | F: AGTCTGTAAGGAAAGCCACCC | Stenhouse et al., 2018 |
|  | R: ACAAAGCCCAAGCAGACAGG |  |
| Pig-  *Lyz1* | F: GGTCTATGATCGGTGCGAGT | Gonzalez et al., 2013 |
|  | R: AACTGCTTTGGGTGTCTTGC |  |
| Pig-  *MUC2* | F: GGCTGCTCATTGAGAGGAGT | Gonzalez et al., 2013 |
|  | R: ATGTTCCCGAACTCCAAGG |  |
| Pig-  *villin* | F: ACGTGTCTGACTCCGAGGGAAAGGT | Yin et al., 2022 |
|  | R: ACTGCTTCGCTTTGATAAAGTTCAG |  |
| Pig-  *Wnt3a* | F: GCGACTTCCTCAAGGACAAG | Gonzalez et al., 2013 |
|  | R: GGTCACGTGTACCGAAGGAT |  |
| Pig-  *β-actin* | F: CACGCCATCCTGCGTCTGGA | Krishna et al., 2015 |
|  | R: AGCACCGTGTTGGCGTAGAG |  |
| Mouse-  *IL-10* | F: GGACCAGCTGGACAACATACTGCTA | Yu et al., 2020 |
|  | R: CCGATAAGGCTTGGCAACCCAAGT |  |
| Mouse-  *TGF-β1* | F: GCTGAACCAAGGAGACGGAAT | Ran et al., 2020 |
|  | R: GCTGATCCCGTTGATTTCCA |  |
| Mouse-  *TNF-α* | F: CCACGCTCTTCTGTCTACTG | Yu et al., 2020 |
|  | R: ACTTGGTGGTTTGCTACGAC |  |
| Mouse-  *IL-1β* | F: GGACAGCCTGTTACTACCTGACACATT | Ding et al., 2020 |
|  | R: CCTAGGAAACAGCAATGGTCGGGAC |  |
| Mouse-  *IL-6* | F: GAGTCACAGAAGGAGTGGCTAAGGA | Yu et al., 2020 |
|  | R: CGCACTAGGTTTGCCGAGTAGATCT |  |
| Mouse-  *IFN-γ* | F: GGACCAGCTGGACAACATACTGCTA | Yu et al., 2020 |
|  | R: CCGATAAGGCTTGGCAACCCAAGT |  |
| Mouse-  *Axin2* | F: AACCTATGCCCGTTTCCTCT | Kim et al., 2021 |
|  | R: GAGTGTAAAGACTTGGTCCA |  |
| Mouse-  *Ctnnb1* | F: ATGGAGCCGGACAGAAAAGC | Kim et al., 2021 |
|  | R: GAGTGTAAAGACTTGGTCCA |  |
| Mouse-  *Hes1* | F: CCAGCCAGTGTCAACACGA | Kim et al., 2021 |
|  | R: AATGCCGGGAGCTACTTTCT |  |
| Mouse-  *Ki67* | F: CCAGCTGCCTGTAGTGTCAA | Sittipo et al., 2020 |
|  | R: TCTTGAGGCTCGCCTTGATG |  |
| Mouse-  *Lgr5* | F: CCTGTCCAGGCTTTCAGAAG | Kim et al., 2021 |
|  | R: CTGTGGAGTCCATCAAAGCA |  |
| Mouse-  *Lyz1* | F: ATGGCGAACACAATGTCAAA | Kim et al., 2021 |
|  | R: GCCCTGTTTCTGCTGAAGTC |  |
| Mouse-  *MUC2* | F: CCTTAGCCAAGGGCTCGGAA | Kim et al., 2021 |
|  | R: GGCCCGAGAGTAGACCTTGG |  |
| Mouse-  *Notch1* | F: GCTGCCTCTTTGATGGCTTCGA | Kim et al., 2021 |
|  | R: CACATTCGGCACTGTTACAGCC |  |
| Mouse-  *Wnt3* | F: CTTCTAATGGAGCCCCACCT | Kim et al., 2021 |
|  | R: GAGGCCAGAGATGTGTACTGC |  |
| Mouse-  *β-actin* | F: TGGAATCCTGTGGCATCCATGAAAC | Kim et al., 2021 |
|  | R: TAAAACGCAGCTCAGTAACAGTCCG |  |

**Table S2.** The information of antibodies used in Immunofluorescence

| Antibody | Company | Catalog | Host species | Dilution |
| --- | --- | --- | --- | --- |
| Pig-β-catenin | Servicebio | GB12016 | mouse | 1:1000 |
| Pig-Ki67 | Servicebio | GB121141 | mouse | 1:300 |
| Pig-Lgr5 | Origene technologies | TA503316 | mouse | 1:100 |
| Pig-MUC2 | Servicebio | GB14110 | mouse | 1:500 |
| Pig-villin | Servicebio | GB121209 | mouse | 1:500 |
| Pig-Wnt3a | Thermo Fisher Scientific | MA-31954 | mouse | 1:200 |
| Mouse-Ki67 | Servicebio | GB111141 | rabbit | 1:500 |
| Mouse-Lyz | Servicebio | GB11345 | rabbit | 1:500 |
| Mouse-MUC2 | Servicebio | GB11344 | rabbit | 1:500 |
| Phalloidine | Servicebio | G1028 |  | 1:300 |
| iFluor^TM^ 594 conjugated Goat anti-mouse IgG | Huabio | HA1126 | mouse | 1:500 |
| Alexa Fluor^®^ 488 conjugated Goat anti-mouse IgG (H+L) | Servicebio | GB25301 | mouse | 1:500 |
| Alexa Fluor^®^ 488 conjugated Goat anti-rabbit IgG (H+L) | Servicebio | GB25303 | rabbit | 1:500 |
| Cy3 conjugated Goat anti-rabbit IgG (H+L) | Servicebio | GB21303 | rabbit | 1:500 |

**Table S3.** The information of antibodies used in Western blot

| Antibody | Company | Catalog | Host Species | Dilution |
| --- | --- | --- | --- | --- |
| Pig-ZO-1 | Thermo Fisher Scientific | 40-2200 | rabbit | 1:500 |
| Pig-Occludin | Abcam | ab222691 | rabbit | 1:500 |
| Pig-Claudin1 | Abcam | ab129119 | rabbit | 1:1000 |
| Pig-E-Cadherin | Thermo Fisher Scientific | 13-1700 | mouse | 1:1000 |
| Pig-β-catenin | Abcam | ab16051 | rabbit | 1:4000 |
| Mouse-Active-β-catenin | Cell Signaling Technology | #19807 | rabbit | 1:1000 |
| Mouse-c-Myc | Cell Signaling Technology | #18583 | rabbit | 1:1000 |
| Mouse-CyclinD1 | Cell Signaling Technology | #55506 | rabbit | 1:2000 |
| Mouse-Lgr5 | Abcam | ab75850 | rabbit | 1:1000 |
| Mouse-Wnt3a | Abcam | ab75850 | rabbit | 1:500 |
| GAPDH | Abcam | ab181602 | rabbit | 1:10000 |
| Goat anti-rabbit IgG (H+L) | Thermo Fisher Scientific | 31210 | rabbit | 1:5000 |
| Goat anti-mouse IgG (H+L) | Thermo Fisher Scientific | 31431 | mouse | 1:5000 |

**Table S4.** The information of antibodies used in Flow cytometry

| Antibody | Company | Catalog |
| --- | --- | --- |
| Fixable Viability Stain 510 | BD Pharmingen | 564406 |
| Ms CD45 PerCP-Cy5.5 30-F11 | BD Pharmingen | 550994 |
| Ms CD3e BV786 145-2C11 | BD Pharmingen | 564379 |
| Ms CD4 FITC RM4-5 | BD Pharmingen | 553046 |
| Ms CD8a APC-Cy7 53-6.7 | BD Pharmingen | 557654 |
| Ms CD25 PE 3C7 | BD Pharmingen | 553075 |
| Ms Foxp3 Alexa 647 MF23 | BD Pharmingen | 560401 |
| Ms ROR Gma T BV650 Q31-378 | BD Pharmingen | 564722 |
| Ms I-A I-E BV605 M5/114.15.2 | BD Pharmingen | 563413 |
| Ms CD11c BV421 HL3 | BD Pharmingen | 562782 |
| Ms CD86 PE-Cy7 GL1 | BD Pharmingen | 560582 |
| Ms CD103 R718 M290 | BD Pharmingen | 752271 |
